# Attenuation of Atherosclerosis with PAR4 Deficiency: Differential Platelet Outcomes in apoE^-/-^ vs. Ldlr^-/-^ Mice

**DOI:** 10.1101/2024.08.01.606266

**Authors:** Caris A. Wadding-Lee, Megan Jay, Shannon M. Jones, Joel Thompson, Deborah A. Howatt, Alan Daugherty, Nigel Mackman, A. Phillip Owens

**Affiliations:** Division of Cardiovascular Health and Disease, Department of Internal Medicine, The University of Cincinnati College of Medicine, Cincinnati, OH, 45267-0542, USA; Pathobiology and Molecular Medicine Program, Department of Internal Medicine, The University of Cincinnati College of Medicine, Cincinnati, OH, 45267-0542, USA; Division of Endocrinology and Molecular Medicine, Department of Internal Medicine, The University of Kentucky Lexington, KY, 40536-0200, USA; Department of Physiology and Saha Cardiovascular Research Center, University of Kentucky, Lexington KY; Division of Hematology, Department of Medicine, UNC Blood Research Center, University of North Carolina at Chapel Hill, Chapel Hill, NC 27599, USA

**Keywords:** Atherosclerosis, protease-activated receptor 4, platelets

## Abstract

**Objective:** Cardiovascular disease (CVD) is a significant burden globally and, despite current therapeutics, remains the leading cause of death. Platelet inhibitors are of interest in CVD treatment to reduce thrombus formation post-plaque rupture as well their contribution to inflammation throughout the progression of atherosclerosis. Protease activated receptor 4 (PAR4) is a receptor highly expressed by platelets, strongly activated by thrombin, and plays a vital role in platelet activation and aggregation. However, the role of PAR4

**Approach and Results:** Mice on a low-density lipoprotein receptor-deficient (*Ldlr*^*-/-*^) background were bred with *Par4* deficient (*Par4*^*-/-*^) mice to create *Ldlr*^*-/-*^*/Par4*^*+/+*^ and *Ldlr*^*-/-*^*/Par4*^*-/-*^ cousin lines. Mice were fed high fat (42%) and cholesterol (0.2%) ‘Western’ diet for 12 weeks for all studies. Bone marrow transplant (BMT) studies were conducted by irradiating *Ldlr*^*-/-*^*/Par4*^*+/+*^ and *Ldlr*^*-/-*^ */Par4*^*-/-*^ mice with 550 rads (2x, 4 hours apart) and then repopulated with *Par4*^*+/+*^ or *Par4*^*-/-*^ bone marrow. To determine if the effects of thrombin were mediated solely by PAR4, the thrombin inhibitor dabigatran was added to the ‘Western’ diet. *Ldlr*^*-/-*^*/Par4*^*-/-*^ given dabigatran did not further decrease their atherosclerotic burden. Differences between apolipoprotein E deficient (*apoE*^*-/-*^) and *Ldlr*^*-/-*^ platelets were assessed for changes in reactivity. We observed higher PAR4 abundance in arteries with atherosclerosis in human and mice versus healthy controls. PAR4 deficiency attenuated atherosclerosis in the aortic sinus and root versus proficient controls. BMT studies demonstrated this effect was due to hematopoietic cells, most likely platelets. PAR4 appeared to be acting independent of PAR1, as there werer no changes with addition of dabigatran to PAR4 deficient mice. *apoE*^*-/-*^ platelets are hyperreactive compared to *Ldlr*^*-/-*^ platelets.

**Conclusions:** Hematopoietic-derived PAR4, most likely platelets, plays a vital role in the development and progression of atherosclerosis. Specific targeting of PAR4 may be a potential therapeutic target for CVD.

**Highlights:**

1. Deficiency of protease-activated receptor 4 attenuates the development of diet-induced atherosclerosis in a *Ldlr*^*-/-*^ mouse model.
2. PAR4 deficiency in hematopoietic cells is atheroprotective.
3. PAR4 deficiency accounts for the majority of thrombin-induced atherosclerosis in a *Ldlr*^*-/-*^ mouse model.
4. The examination of platelet-specific proteins and platelet activation should be carefully considered before using the *apoE*^*-/-*^ or *Ldlr*^*-/-*^ mouse models of atherosclerosis.

## INTRODUCTION

Coronary artery disease (CAD) is a common cause of cardiovascular disease and death globally, accounting for approximately 1M deaths every year in the US.^1^ Atherosclerosis, is the underlying cause of CAD, is chronic inflammatory condition marked by subendothelial accumulation of lipoproteins, immune cells, and vascular smooth muscle cells (VSMCs) within artery walls.^2^ Critical to initiation of atherosclerosis, platelets affix to active endothelium shedding inflammatory chemokines like C-C motif chemokine ligand 5 (CCL5) and C-X-C motif chemokine ligand 12 (CXCL12), luring circulating leukocytes to stick firmly to vessel wall and infiltrate into lesions.^3^ Moreover, platelets release inflammatory cytokines, such as interleukins and CD40 ligand, hastening endothelial cell activation, boosting adhesion molecule production and display, and further accelerating atherosclerosis. Even though statins and antithrombotic therapies targeting platelet ADP receptors and thromboxane pathways are used widely, the risk for recurrent atherothrombosis and adverse coronary events three years after percutaneous coronary intervention (PCI) remains high at ∼20%.^4^ This showcases the urgent demand for new treatments halting or reversing atherosclerotic lesions progress without compromising safety.

Platelets play an extensive role in development and progression of atherosclerosis. Atherosclerosis in high fat and cholesterol diet-fed low-density lipoprotein deficient mice (*Ldlr*^*-/-*^) is blunted by platelet depletion due to reduced monocyte chemotaxis and macrophage accumulation.^5^ Thrombin serves as a potent activator of platelets by engaging protease-activated receptors (PARs). The four known PARs (PAR1, PAR2, PAR3, and PAR4) are activated through cleavage of their N-terminal domains by serine proteases, which exposes a tethered ligand for receptor activation.^6^ Notably, human platelets express PAR1 and PAR4 as receptors for thrombin.^7^ While initial studies have pointed to PAR1 as the main receptor involved in thrombin-induced platelet activation, recent research outlines distinct roles for both PAR1 and PAR4.^8,9^ Development of the PAR1 antagonist, vorapaxar, has shown clinical effectiveness but also increases bleeding risks, which has generated interest in PAR4 antagonists as an alternative antithrombotic agent.^10^ In mice, thrombin-induced platelet activation is mediated by PAR4 with PAR3 acting as a cofactor.^11^ Studies utilizing direct thrombin inhibitors or prothrombotic mouse models indicate a role of thrombin in the onset of atherosclerosis. This underscores the multifaceted roles of thrombin and its receptors in vascular biology.

We demonstrated previously that PAR1 does not play a significant role in development or progression of atherosclerosis in *Ldlr*^*-/-*^ mice.^2^ While this effect would likely be platelet-independent, as PAR1 is not expressed in mouse platelets, another study determined PAR1 on a apolipoprotein E deficient (*apoE*^*-/-*^) background was protected from diet-induced atherosclerosis due to decreased cholesterol efflux and leukocyte recruitment.^12^ Interestingly, two studies concluded that PAR4 does not affect atherosclerosis in an *apoE*^*-/-*^ mouse model or in commonly characterized mouse atherosclerotic vessels (aortic sinus and *en face* aorta) in a scavenger receptor B1 (*Sr-b1)/Ldlr*^*-/-*^ mouse using a PAR4 pepducin inhibitor RAG8.^13,14^ However, the role of thrombin in initiation and progression of atherothrombosis-induced CAD is indisputable. The *in vivo* generation of thrombin, as measured by thrombin-antithrombin complexes (TAT) or prothrombin fragment 1.2 (F1.2) are both acutely and chronically increased in patients with symptomatic CAD.^15-18^ Moreover, these thrombin-generated products remain elevated for up to two years following an event and occur again for patients with restenosis.^19,20^ This increase in thrombin generation products is correlated with the severity of calcification and overall arterial disease.^21^ Studies link initial plasma F1.2 concentrations to the risk of sudden death or re-infarction post an acute myocardial infarction (MI). Further, elevated plasma TAT concentrations to negative outcomes in ST-segment elevation myocardial infarction (STEMI) patients.^22,23^ Recombinant thrombomodulin (TM) rTMD23 results in augmented thrombin binding to TM and reduces atherosclerosis and neointima formation in a *apoE*^*-/-*^ mouse model.^24^ Alternatively, pro-thrombotic *TM*^*Pro/Pro*^ mice, with decreased TM-binding to thrombin, have severe atherosclerosis, increased lesion vulnerability, and spontaneous atherothrombosis in *apoE*^*-/-*^ mice, which is rescued with the thrombin inhibitor dabigatran or administration of recombinant activated protein C (APC).^25^ Thrombin is considered one of the most potent agonists for platelet activation,^26^ but the link between thrombin and platelets in mouse models of atherosclerosis remains understudied. Therefore, while the link between thrombin activity and CAD along with its issues is clear, the depth and specifics of these connections to platelets requires further exploration.

## MATERIALS AND METHODS

### Mice and diet

*Par4*^*-/-*^ mice were obtained from Dr. Coughlin (UCSF) and were 6 times backcrossed into C57BL/6J, respectively.^27^ *Ldlr*^*-/-*^ male mice were obtained from The Jackson Laboratory. *Ldlr*^*-/-*^ */Par4*^*+/+*^ and *Ldlr*^*-/-*^*/Par4*^*-/-*^ cousin lines were generated by interbreeding *Ldlr*^*-/-*^*/Par4*^*-/-*^ onto the *Ldlr*^*-/-*^ strain. *apoE*^*-/-*^ mice for platelet studies were obtained from The Jackson Laboratory.

All mice were provided normal mouse laboratory diet and water ad libitum. To induce hypercholesterolemia, mice were fed a ‘Western’ diet (TD.88137). For inhibition of thrombin, mice were fed a custom-made ‘Western’ diet containing peanut butter (10g/kg diet) with or without dabigatran (10g/kg diet).

### Bone marrow transplantation (BMT)

BMT was conducted as previously described.^2^

### Plasma collection and processing of heart and aorta

Plasma was collected and the heart and aorta processed as described previously.^2^

### Aortic sinus atherosclerosis quantification

Atherosclerotic lesions in the aortic sinus were sectioned, stained and analyzed as described previously.^28,29^

### En face atherosclerosis quantification

En face atherosclerosis was performed and quantified, as described previously.^28,29^

### Immunostaining for CD68

Immunostaining was performed on frozen serial sections as described previously.^30^

### Quantitative real-time polymerase chain reaction (qRT-PCR)

qRT-PCR was performed as described previously.^2^ Taqman probes listed in Supplemental Material.

### Human carotid artery atherosclerotic samples

Human carotid artery atherosclerotic samples were obtained from Origene Technologies, Inc. and processed as described previously.^2^

### PAR4 ELISA

PAR4 protein in both mouse and human samples were measured utilizing a sandwich ELISA.

### Platelet isolation

Platelets were collected and washed as described previously.^31^

### Microtiter plate-based light transmission aggregometry

All incubations and assays were performed as described previously.^31^

### Research statistics and data representation

All statistical analysis and graphical output was generated using SigmaPlot v.15 (SPSS, Chicago, IL), or GraphPad (v10.2.3, GraphPad software LLC). Data are represented as mean ± SEM. For two group comparison of parametric data, a Student’s t-test was performed, while non-parametric data was analyzed with a Mann-Whitney Rank Sum. Statistical significance between multiple groups was assessed by One Way analysis of variance (ANOVA) on Ranks with a Dunn’s post hoc, One Way ANOVA with Holm Sidak Post Hoc, or Two-Way ANOVA with Holm Sidak Post Hoc, when appropriate. Statistical significance among groups performed temporally was assessed by either a One-Way Repeated Measures ANOVA or Repeated Measures ANOVA on Ranks, where appropriate. Values of *P* < 0.05 were considered statistically significant.

### Study approvals

All mouse studies were performed with the approval of the University of North Carolina at Chapel Hill and the University of Cincinnati Institutional Animal Care and Use Committees.

## RESULTS

### PAR4 abundance was increased in mouse and human atherosclerotic lesions

PAR4 abundance was measured in normal and atherosclerotic vessels of mice and humans. Male *Ldlr* ^*-/-*^ mice were fed either a control laboratory diet or a high fat and cholesterol “Western” diet for 24 weeks prior to harvesting the aortic arches. Human carotid artery samples with or without atherosclerosis were used to examine PAR4 abundance in humans. These samples were age-matched human arteries between atherosclerotic burdened carotid arteries and controls. We found increased *Par4* mRNA and protein abundance in both atherosclerotic mouse models as well as human atherosclerosis burdened carotid arteries versus their healthy controls (Figures 1 A-D).

**Figure 1:**
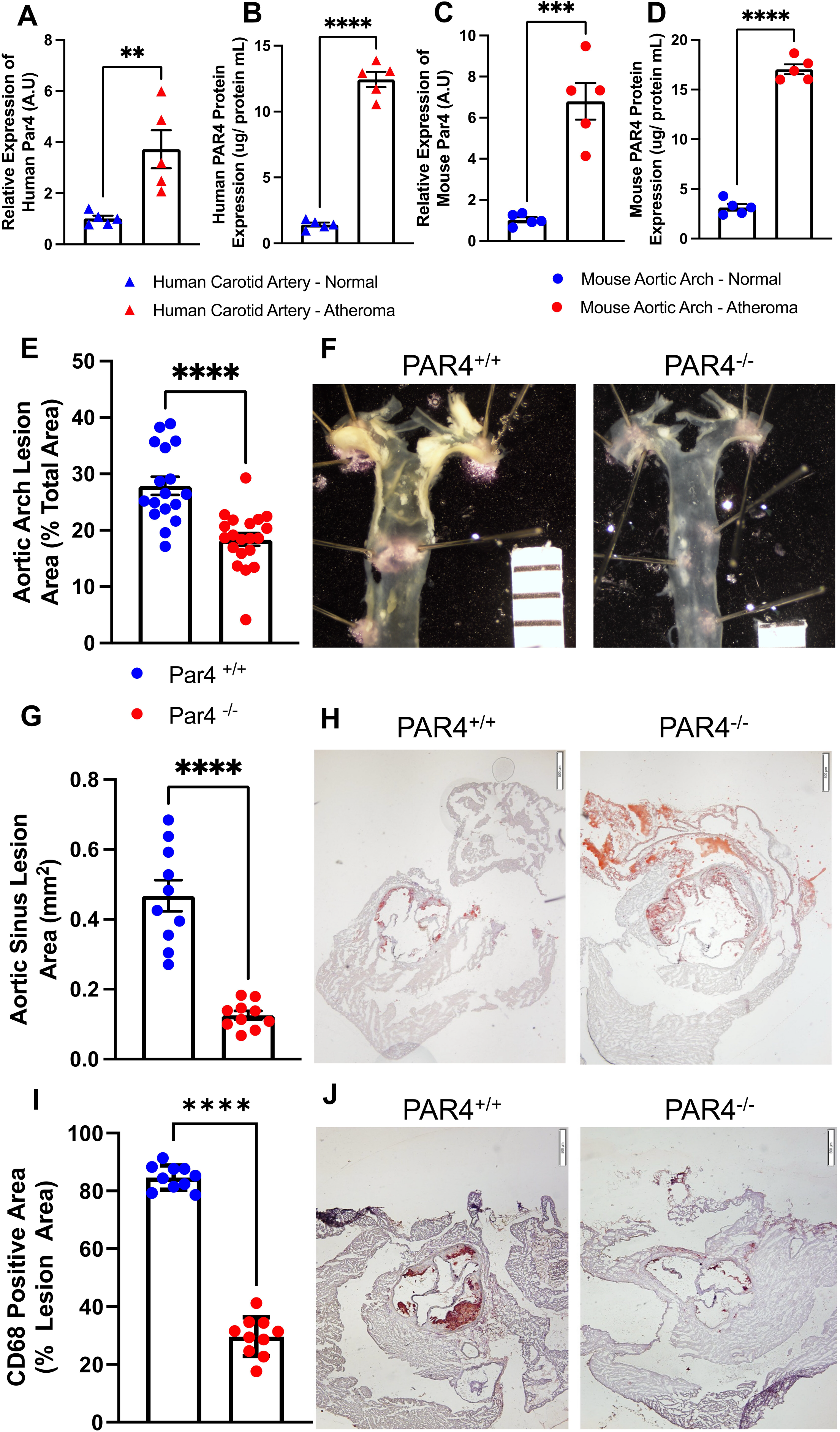
PAR4 deficiency reduced formation of atherosclerosis in *Ldlr*^*-/-*^ mice. **A-D:** Normal mouse aortic arch (*Ldlr*^*-/-*^ mice fed chow diet for 24 weeks) or atherosclerosis-induced aortic arches (*Ldlr*^*-/-*^ mice fed a Western diet for 24 weeks) were examined for (A) Par4 mRNA expression normalized to HPRT and (B) PAR4 protein expression (n=5 individual samples/ group). Normal or atheroma burdened human carotid arteries were isolated and examined for (C) *Par4* mRNA expression normalized to 18s and (D) PAR4 protein expression (n=5 individual samples/ group). Histobars represent mean + SEM, triangles represent individual human values, and circles represent individual mouse values. Student t-test were completed for each graph where **denotes P=0.0069, ***denotes P=0.0002, and ****denotes P<0.0001 comparing atheroma samples are compared to normal artery condition. **E-J:** 8-12-week male mice on a *Ldlr*^*-/-*^ background that were *Par4*^*+/+*^ or *Par4*^*-/-*^ (n=10-20) were fed a high fat/ cholesterol diet for 12 weeks. (E) Percent lesion area of the aortic arch and (F) representative images of unstained en face aortic arches from *Par4*^*+/+*^ and *Par4*^*-/-*^ mice. (G) Quantification of aortic sinus with lesion area and (H) Oil-red O stained aortic sinuses from *Par4*^*+/+*^ and *Par4*^*-/-*^ mice. (I) Quantification of CD68 stain represented in (J) aortic sinuses sections. Individual measurements are represented in circles and histobars represent means +/-SEM. ****Denotes P<0.0001 comparing deficient to proficient mice using Student’s t-test.

### PAR4 deficiency was associated with attenuation of diet-induced atherosclerosis in mice

To investigate the role of PAR4 in atherosclerotic development, we used 8 – 12-week-old male mice *Ldlr*^*-/-*^*/Par4*^*+/+*^ and *Ldlr*^*-/-*^*/Par4*^*-/-*^ (n = 10 – 20/ genotype) that were fed a Western diet for 12 weeks. *Ldlr*^*-/-*^*/Par4*^*-/-*^ mice showed significantly less atherosclerotic development in the aortic arch and the aortic sinus compared to controls (Figures 1E–H). Histological sections were analyzed to determine macrophage accumulation within the lesion area of these mice. *Par4* deficient mice had less macrophage accumulation compared to proficient controls (Figure 1I and 1J). Body weight comparisons completed throughout the study show more body weight gain in PAR4 deficient mice versus proficient controls (Supplemental Figure IA), though initial weights did not show a difference between PAR4 deficient and proficient mice (Supplemental Figure IB). Analyses of fat depots showed larger epididymal and retroperitoneal fat pads in PAR4 deficient mice as well as larger livers (Supplemental Figures IC and ID).

### PAR4 expressed by hematopoietic cell type contributed to atherosclerotic development

We used bone marrow transplant to determine the cellular source of PAR4 on atherosclerosis. *Ldlr*^*-/-*^*/Par4*^*+/+*^ and *Ldlr*^*-/-*^*/Par4*^*-/-*^ male mice (8 – 10 weeks old, n = 20/genotype) were irradiated and then transplanted with bone marrow cells from either *Ldlr*^*-/-*^*/Par4*^*+/+*^ or *Ldlr*^*-/-*^*/Par4*^*-/-*^ to create a total of 4 chimeric groups. After a 5-week interval to allow for bone marrow repopulation, mice were fed a ‘Western’ diet for 12 weeks to induce atherosclerosis. Body weights collected throughout the Western diet study show no difference in weight gain between the different groups and no difference in initial weights prior to starting diet (Supplemental Figures IIA and IIB). Mice repopulated with Par4 deficient bone marrow had less atherosclerotic lesion development versus mice repopulated with Par4 proficient bone marrow (Figures 2A – D). These results were seen in both the aortic arch as well as the aortic sinus. Mice with *Par4* deficient bone marrow had less macrophage accumulation in their atherosclerotic lesions versus mice repopulated with *Par4* proficient bone marrow (Supplemental Figure IIC and IID).

**Figure 2:**
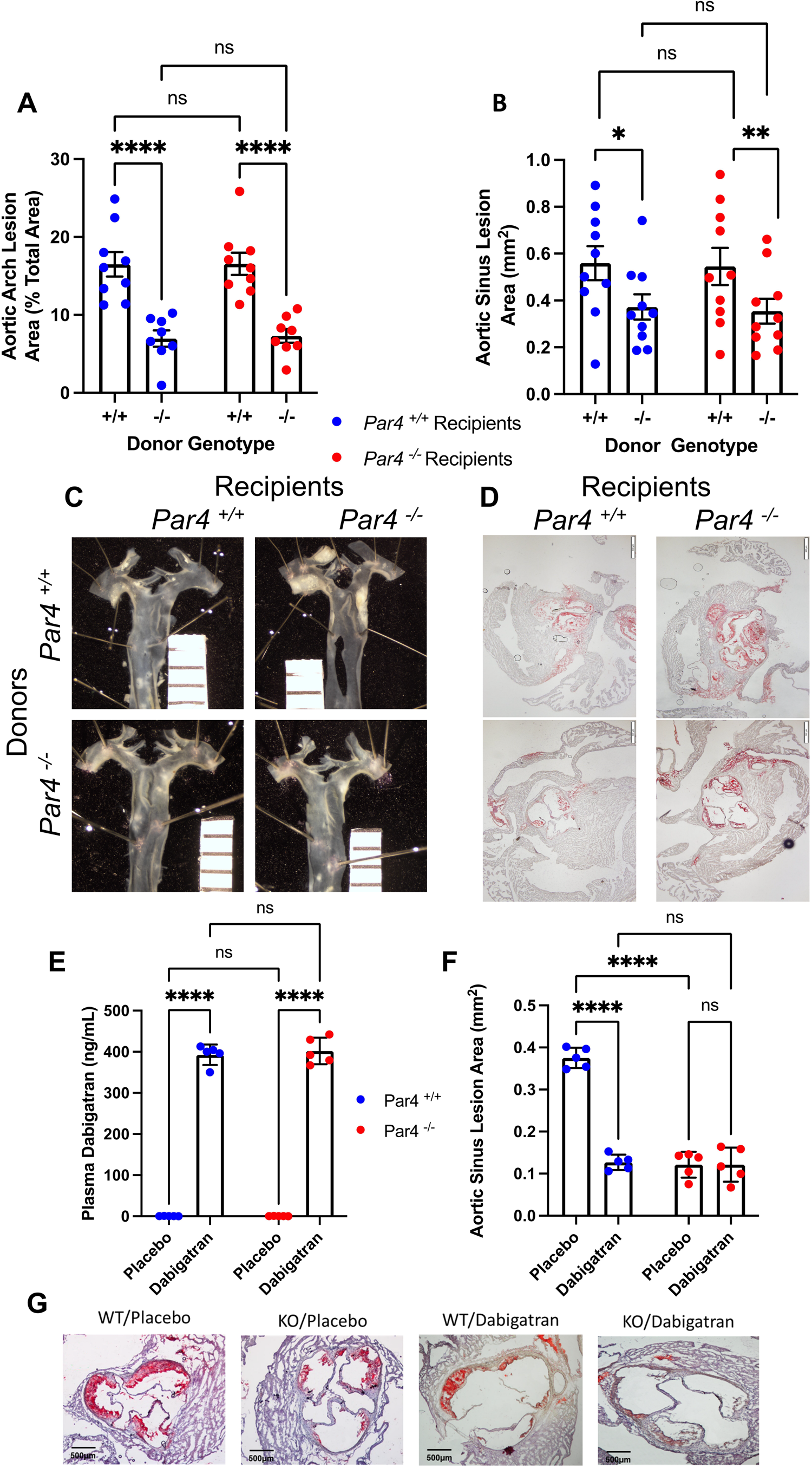
Hematopoietic PAR4 reduced atherosclerosis in *Ldlr*^*-/-*^ mice. **A-D:** Male *Ldlr*^*-/-*^ /*Par4*^*+/+*^ and *Ldlr*^*-/-*^*/Par4*^*-/-*^ (8-12 weeks old, n=20/group) were irradiated of their bone marrow using two rounds of 550 rads of radiation. Each genotype were split into two “recipient” groups, either repopulated with *Par4*^*+/+*^ or *Par4*^*-/-*^ bone marrow, given reto-orbitally. Mice that were used for bone marrow collections are termed “donor”. Mice were rested for a period of four weeks and then fed Western diet for 12 weeks. (A) Percent lesion area of unstained aortic arch atherosclerosis and (B) representative images of unstained *en face* aortic arches. (C) Quantified aortic sinus lesion area and (D) Oil-red O stained aortic sinuses. Individual measurements are represented with circles and histobars represent mean +/-SEM. Two-way ANOVA with Multiple Comparisons where ****denotes P<0.0001, **denotes P=0.0463, and *denotes P=0.05. **E-G:** Male *Ldlr*^*-/-*^ mice (8-12 weeks) that were either *Par4*^*+/+*^ or *Par4*^*-/-*^ (n=5/ genotype) were fed 10mg/kg of dabigatran diet on a Western background or fed placebo diet. These mice were fed diet for a period of 12 weeks. (A) Represents plasma dabigatran levels in ng/mL and (B) aortic sinus lesion area where circles represent individual measurements and histobars represent means +/-SEM. Two Way ANOVA with multiple comparisons was done where ****denotes P<0.0001.

### Thrombin inhibition did not further attenuate atherosclerosis beyond PAR4 deficiency

PAR4 is the sole thrombin receptor on mouse platelets and is known to augment the development and progression of various cardiovascular diseases.^11,32^ To determine if PAR4 is contributing to the majority of thrombin-induced effects in our atherosclerotic mouse models, we used *Ldlr*^*-/-*^*/Par4*^*+/+*^ and *Ldlr*^*-/-*^*/Par4*^*-/-*^ male mice (8 – 12 weeks old, n = 5/genotype) that were fed either a ‘Western’ diet mixed with a thrombin inhibitor dabigatran (10 mg/kg) or cellulose-based placebo ‘Western’ diet. Mice were fed these diets for a period of 12 weeks and humanly euthanized for analysis of atherosclerotic lesion size in the aortic sinus. Plasma dabigatran concentrations were measured in all mice. Mice fed dabigatran had significantly increased dabigatran in plasma, while dabigatran was undetectable in the plasma of placebo-fed mice (Figure 2A). Body weights collected through the diet study show no difference in weight gain between the different groups as well as no significant difference in body weight prior to starting the diet study (Supplemental Figure IIIA and IIIB). Administration of dabigatran significantly reduced aortic sinus atherosclerosis in *Par4+/+* mice compared to placebo controls (Figure 2F – 2G). Interestingly, dabigatran did not reduce the amount of atherosclerotic burden lower than *Par4* -/- alone (Figure 2).

### ApoE deficiency augmented platelet reactivity compared to Ldlr deficient mice

At present, it is not clear why our results differ from Hamilton and colleagues.^14^ The one difference between our two studies is Hamilton’s use of an *apoE*^*-/-*^ mouse model, while our studies were conducted on a *Ldlr*^*-/-*^ background strain. Previous studies have shown that *apoE* -/- mice have altered platelet activation, as apoE inhibits platelet activation via nitric oxide-induced pathways.^33^ To examine the differences in platelet activation between these two mouse strains, male *apoE*^*-/-*^ and *Ldlr*^*-/-*^ mice platelets were harvested at baseline (day 0; n = 8 each) and after 12 weeks of ‘Western’ diet (day 84, n = 8 each). At baseline, there were no differences between *apoE*^*-/-*^ and *Ldlr*^*-/-*^ mouse platelets incubated with selected concentrations of the agonists thrombin (PAR4 agonist) and collagen (glycoprotein VI – GPVI agonist; Figure 5). *Ldlr*^*-/-*^ mice fed a ‘Western’ diet had a significant increase over *apoE*^*-/-*^ and *Ldlr*^*-/-*^ baseline, while the platelets of ‘Western’ fed *apoE*^*-/-*^ mice were significantly increased over all other groups (Supplemental Figure IV). Platelets derived from our study mice *Ldlr*^*-/-*^*/Par4*^*+/+*^ and *Ldlr*^*-/-*^*/Par4*^*-/-*^ (fed ‘Western’ for 12 weeks) were similar to platelets from Western-fed Ldlr^-/-^ treated with collagen. Thrombin-mediated effects were similar for *Ldlr*^*-/-*^*/Par4*^*+/+*^ mouse platelets, but completely ablated in *Ldlr*^*-/-*^*/Par4*^*-/-*^ mice (Supplemental Figure V).

## DISCUSSION

The results of our study examining *Par4*^*-/-*^ in *Ldlr*^*-/-*^ mice significantly differ from a previous examination of PAR4 deficiency in *apoE*^*-/-*^ mice.^14^ As such, it is important to note the potential impacts of *apoE* versus *Ldlr* deficient mouse models can have on platelet behavior. Platelets express the receptor apoE receptor 2 (*apoER2*) and low-density lipoprotein receptor-protein 8 (LRP8) that binds and internalizes apoE.^34,35^ This action of this receptor is independent of lipid uptake and resultant binding of apoE to LRP8 leads to decreased platelet activation and aggregation.^35,36^ Further suggesting a role of apoE in platelet activation and aggregation, *Lrp8* deficiency results in reduced ADP and thrombin-induced platelet activation and increased time for carotid occlusion.^36^ Importantly, apoE inhibition of platelet reactivity was attenuated in *Lrp8* deficient platelets, although some inhibition remained, indicating the potency of apoE as an anti-platelet agent and indicating additional non-LRP8 pathways.^33,36^ As apoE plays an imperative role in platelet homeostasis and platelet activation in disease states, ablating apoE could potentially alter the effects of platelet response in an atherogenic mouse model. Furthermore, platelets interact with macrophages to influence apoE expression as well as augmenting macrophage foam cell formation.^37,38^ Conversely, Ldlr is not expressed on platelets, and exogenous addition of LDL does not influence platelet activation, thus deletion of Ldlr should not have an impact on platelet studies in atherosclerotic mouse models.^34,36^

In addition to receptor expression on platelets, *apoE*^*-/-*^ and *Ldlr*^*-/-*^ atherosclerotic mouse models have differences that should be taken into consideration. The lipoprotein profiles of *apoE*^*-/-*^ and *Ldlr*^*-/-*^ mice differ, with *apoE*^*-/-*^ mice showing a higher concentration of chylomicrons and VLDL remnants (mainly apoB-48), whereas *Ldlr*^*-/-*^ mice had higher concentrations of LDL (mainly apoB-100).^39^ Importantly, apoB-100 plays a significant role in lipoproteins’ ability to bind proteoglycans expressed by the vascular surface and enter the subendothelial space to assist in the initiation and progression of atherosclerosis.^40^ When both *apoE*^*-/-*^ and *Ldlr*^*-/-*^ mice are genetically altered to express only apoB-100, *Ldlr*^*-/-*^ mice display higher LDL lipid profiles and augmented en face aortic atherosclerosis versus *apoE*^*-/-*^ mice with a VLDL-dominant lipid profile, suggesting *Ldlr*^*-/-*^ mice are able to accumulate lipoproteins more readily in the intima than *apoE*^*-/-*^ mice.^41^ An in-depth examination of *apoE*^*-/-*^ and *Ldlr*^*-/-*^mice demonstrate both strains show a prothrombotic phenotype due to ensuing hypercholesterolemia with both distinctive platelet characteristics and lipid profiles.^42^ *ApoE*^*-/-*^ mice have increased platelet aggregation with moderate increases in thrombin potential, whereas *Ldlr*^*-/-*^ mice display increase platelet secretion without changes in plasma thrombin.^42^ While it was concluded that both mouse strains had increased thrombotic processes due to increases in cholesterol-induced platelet activation, *apoE*^*-/-*^ mouse platelets also had increased inflammatory lipids.^42^ Similarly, our data demonstrates there are no differences in the platelet reactivity between normal laboratory diet fed *apoE*^*-/-*^ and *Ldlr*^*-/-*^ mice. However, platelets isolated from ‘Western’ diet-fed *Ldlr*^*-/-*^ mice have increased reactivity to thrombin and collagen versus both normal laboratory diet fed groups. Additionally, platelets isolated from ‘Western’ diet-fed *apoE*^*-/-*^ mice have increased reactivity versus all other groups. Together, this data suggests caution in both selecting and interpreting studies with platelet-derived proteins and platelet activation/secretion in mouse models of atherosclerosis.

Similar to our studies, other investigators have found dabigatran reduces lesion area in mouse models of atherosclerosis, with mechanisms ranging from reductions in oxidative stress and inflammation.^43^ While the PAR1 antagonist, vorapaxar, has clinical effectiveness, it has also been associated with increased bleeding risks, and there is a general hesitancy to move away from P2Y12 inhibitors and aspirin or to move to ‘triple’ platelet therapy.^10^ Recently, interest has been generated toward PAR4 antagonists, as PAR4 has a higher action potential for activation by thrombin and may result in less bleeding risk with PAR1 as a redundant thrombin receptor. A recent study utilizing a pepducin inhibitor of PAR4 (RAG8) found that when platelet activation was decreased, there was less platelet accumulation in coronary atherosclerotic lesions, and coronary lesion area was decreased in scavenger receptor class B type I (SR-B1) deficient mice on a *Ldlr*^*-/-*^ background.^13^ Interestingly, the atherosclerotic burden in the ‘normal’ vascular beds of the aortic sinus and *en face* aortic root were not changed with the PAR4 inhibitor.^13^ The selective oral PAR4 antagonist, at a dose of 60 mg, inhibited PAR4 activation and subsequent platelet aggregation and activation for a short period (two hours), which then returned to baseline levels after 24 hours.^44^ Thrombus formation was also reduced versus previously published doses of aspirin and clopidogrel.^44^ A cohort of patients treated with PAR4 antagonist BMS-986141 demonstrated a reduction of thrombus formation patients with stable CHD, hypothesized to be due to a inhibition of PAR4’s ability to activate platelets for a prolonged period of time when activated via thrombin and inhibiting this action in turn decreases the platelet presence within the thrombus itself.^45^

Our study has shown that targeting PAR4 in models of CAD could be a viable therapeutic for decreasing the burden this disease holds on the global population. In addition to several other mouse studies, we have shown that deletion of PAR4 in a *Ldlr*^*-/-*^ atherosclerotic mouse model decreases lesion area as well as macrophage accumulation within the vasculature. We have determined that these effects are due to a hematopoietic cell type, with high potential it is platelet expressed PAR4 causing the progression of atherosclerosis. As shown by several clinical and preclinical studies, thrombin inhibition and, even more so, PAR4 inhibition are viable therapeutics that reduce cardiovascular health risk and have few side effects in patients.

## Acknowledgments

We thank the countless undergraduate students who have worked on this project throughout the years. Thanks to Kellie Machlus, Ph.D. and Martina Slingsby at Boston Children’s/Harvard for her invaluable feedback regarding the microtiter plate assay.

## Nonstandard Abbreviations and Acronyms

APC: Activated protein C
apoE: Apolipoprotein E
BMT: Bone marrow transplant
CAD: Coronary artery disease
DTI: Direct thrombin inhibitors
LDL: Low density lipoprotein
PAR: Protease activated receptor
PCI: Percutaneous coronary intervention
STEMI: ST-segment elevation myocardial infarction
TAT: Thrombin antithrombin
TM: Thrombomodulin
VCAM1: Vascular cell adhesion molecule 1

